# α-Synuclein Facilitates Spontaneous Dopamine Release in a Calcium- and Phosphorylation-Dependent Manner

**DOI:** 10.64898/2026.04.20.719002

**Authors:** Yuqing Feng, Amberley D. Stephens, Pedro Vallejo Ramirez, Eugene V. Mosharov, Alfonso De Simone, Giuliana Fusco, Stanislaw Makarchuk, Marius Brockhoff, Ana Fernandez-Villegas, Colin Hockings, Edward Ward, Pedro Magalhães, Senthil Kumar, Nino F. Läubli, Hilal A. Lashuel, Clemens F. Kaminski, Gabriele S. Kaminski Schierle

## Abstract

α-Synuclein (aSyn) is central to Parkinson’s disease pathogenesis, yet its native physiological role at the presynapse remain poorly defined. Here, super-resolution imaging in dopaminergic neurons reveals that endogenous aSyn localises within nanometres of L-type voltage-gated calcium channels (LTCC), with closer proximity under both spontaneous neuronal activity and stimulated conditions compared to when extracellular calcium is chelated. Blocking Ca^2+^/calmodulin-dependent kinase II (CaMKII) reduces aSyn clustering at LTCC under spontaneous activity, suggesting that calcium entry and downstream calcium-dependent kinase activity contribute to aSyn localisation. Moreover, quantitative single-molecule analyses indicate that calcium increases the abundance of both total and serine129 phosphorylated (pS129) aSyn in synaptosomes under spontaneous conditions, and NMR analysis reveals that both calcium and S129 phosphorylation increase the binding affinity of aSyn to synaptic vesicles. Functional assays further demonstrate that LTCC blockade elevates intracellular DA levels exclusively in the presence of aSyn under spontaneous but not stimulated conditions. Finally, biochemical fractionation and multi-colour single-molecule imaging reveal that aSyn preferentially associates with small vesicles that are not obligately coupled to full-fusion associated recycling pools. These results suggest that aSyn acts as a calcium- and phosphorylation-regulated modulator of spontaneous DA release through pathways that are largely independent of full-fusion recycling mechanisms, and that pS129 aSyn is not solely a pathological marker but may also reflects physiological regulation. Together, these insights provide a framework for understanding how therapeutic strategies targeting aSyn may impact its normal synaptic functions.

## Introduction

Despite nearly four decades of research linking α-synuclein (aSyn) to the pathogenesis of Parkinson’s disease (PD), its presynaptic organisation, interactions, and functions in the native cellular environment remain poorly defined. A clearer understanding of these physiological properties is critical for evaluating emerging therapeutic strategies that aim to modulate or reduce aSyn levels. However, many models of aSyn function have relied on overexpression systems, which can profoundly distort its localisation and activity (1,2), thus complicating the interpretation of its native roles.

Structurally, aSyn is a 140-amino-acid intrinsically disordered protein (IDP) comprising three principal domains: an amphipathic N-terminal region that adopts an α-helical conformation upon membrane binding (3); a hydrophobic non-amyloid-β component (NAC) central region that drives aggregation (4); and an acidic C-terminal region that imparts chaperone-like activity and hosts most post-translational modifications (PTMs) (5,6). The C-terminus exhibits weak intrinsic membrane affinity but can regulate membrane-bound interactions and protein-protein contacts (7,8). Recent studies have revealed that calcium binds to aSyn’s C-terminus (9), potentiating its interaction with synaptic vesicles (10). Furthermore, phosphorylation at S129 has long been associated with aSyn pathology, but there is ongoing debate about whether it contributes to aSyn toxicity or modulates its normal functions (11,12). Thus, elucidating the interplay between calcium interactions and post translation modification (PTM) of aSyn is critical, not only to understand its physiological function(s) at synaptic terminals, but also to anticipate the consequences of therapeutic interventions targeting aSyn.

aSyn has been proposed to regulate vesicle organisation and trafficking at presynaptic terminals (13), contributing to vesicle clustering and docking (14–16), and promoting soluble N-ethylmaleimide-sensitive factor attachment protein receptor (SNARE) complex assembly to modulate neurotransmitter release (17). In addition to exocytosis, aSyn has also been implicated in vesicle recycling via non-canonical fusion and endocytic pathways (1,18). Despite these proposed functions, how aSyn activity is regulated by neuronal activity and calcium signalling, and whether it acts on specific vesicle populations under physiological (i.e., not overexpressed) conditions remains incompletely defined.

In this study, we address this gap by exploring how extracellular calcium and neuronal activity influence the spatial localisation and phosphorylation of endogenous aSyn, and how these changes relate to its interactions with synaptic vesicles and regulation of dopamine (DA) release. Using DNA points accumulation for imaging in nanoscale topography (DNA-PAINT), a super-resolution imaging method, together with single-molecule quantification (qPAINT), and vesicle biochemistry, we characterise whether aSyn recruitment, phosphorylation state, and vesicle-binding affinity depend on calcium availability. Complementary functional assays and synaptosome direct stochastic optical reconstruction microscopy (dSTORM) imaging further clarify whether aSyn engages canonical full-fusion associated recycling or partakes in alternate DA release pathways.

Together, our results reveal that calcium is essential for presynaptic enrichment of aSyn in the vicinity of LTCCs and for vesicle-binding behaviour, and that spontaneous neuronal activity dynamically regulates its phosphorylation. Additionally, aSyn is more associated with small vesicles and supports DA release largely independent of full-fusion associated recycling. These findings provide a critical mechanistic framework for understanding how calcium- and phosphorylation-dependent regulation of aSyn interactions may shape both its physiological roles and the effectiveness of therapeutic strategies aimed at its clearance or stabilisation.

## Results and Discussion

### Calcium and CaMKII activity increase the proximity of aSyn to LTCCs under spontaneous activity

We have previously shown that α-synuclein (aSyn) binds multiple Ca^2+^ ions at its C-terminus and interacts with synaptic vesicles in a calcium-dependent manner (10). This suggested that aSyn may be recruited to sites of elevated calcium flux, particularly in the vicinity of voltage-gated calcium channels (VGCCs) where calcium concentrations reach the peak during neuronal activity. In this study, we focused on L-type VGCCs (LTCCs) because of their established role in dopaminergic neurons pacemaking activity (19), their contribution to the vulnerability of substantia nigra neurons in Parkinson’s disease (PD) (20),and the protective effects of LTCC inhibitors observed in preclinical PD models (21). Additionally, recent evidence indicates that aSyn and LTCC are functionally connected (22,23). To provide structural evidence for this interaction, we employed DNA-PAINT super-resolution microscopy to map the nanoscale spatial relationship between aSyn and LTCCs (targeting Ca_v_1.3 channels) in dopaminergic neuronal cultures (Fig. 1a).

**Figure 1.**
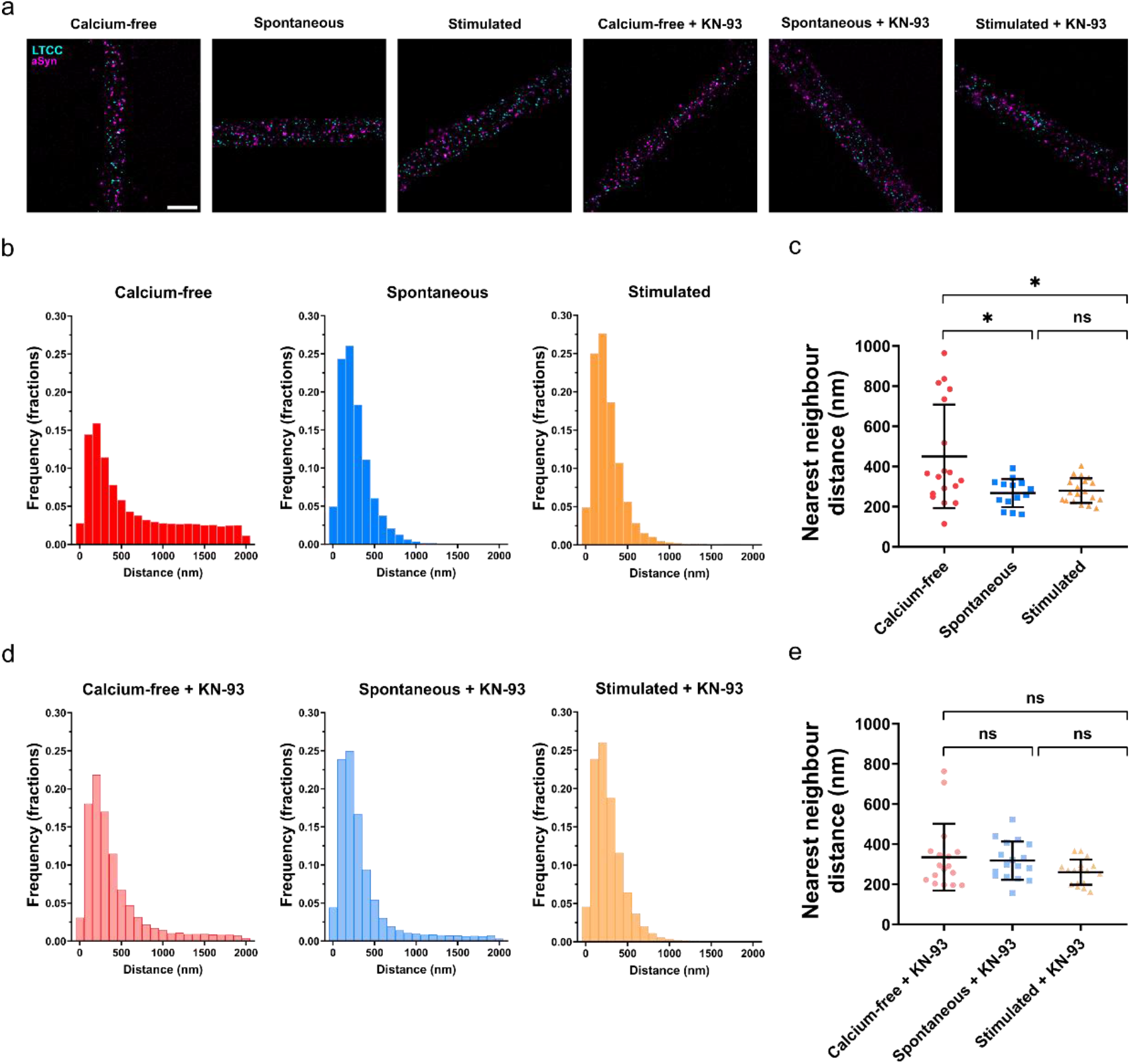
Calcium and CaMKII activity promote aSyn’s proximity to LTCCs. **(a)** Representative reconstructed DNA-PAINT images of aSyn (magenta) and LTCCs (cyan) in neurites of P2 rat dopaminergic neurons under calcium-free (1 mM EGTA), spontaneous (2 mM CaCl_2_), or stimulated conditions (70 mM KCl and 2 mM CaCl_2_), with or without KN-93 (a CaMKII inhibitor). Scale bar, 2 µm. **(b)** Histograms of nearest-neighbour distances from aSyn to the closest LTCC (bin width, 100 nm). **(c)** Mean ± s.d. of nearest-neighbour distances per image in the three buffer conditions. Brown-Forsythe and Welch ANOVA with Tamhane’s T2 multiple comparisons. ns, not significant; *, P < 0.05. Data from N = 3 independent biological repeats were quantified. **(d)** Histograms of nearest-neighbour distances from aSyn to the closest LTCC for conditions with KN-93 treatment (bin width, 100 nm). **(e)** Mean ± s.d. of nearest-neighbour distances per image across buffer with KN-93 treatment. Kruskal-Wallis with Dunn’s multiple comparisons. ns, not significant. Data from N = 3 independent biological repeats were quantified.

Postnatal day 2 (P2) rat primary dopaminergic neurons were incubated in extracellular buffer under three conditions: calcium-free (1 mM EGTA), spontaneous (resting), allowing tonic neurotransmitter release (2 mM CaCl_2_) and stimulated, inducing evoked neurotransmitter release (70 mM KCl and 2 mM CaCl_2_). Under calcium-free conditions, nearest-neighbour analysis revealed that more aSyn molecules were detected within 0-2 µm from LTCCs (average number of pairs per image: 2290), with distribution across the entire 0-2 µm range (Fig. 1b). In contrast, spontaneous and stimulated conditions had fewer aSyn molecules within 0-2 µm from LTCCs (physiological: 1345, stimulated: 1153) but a significantly higher proportion localised within ~300 nm, leading to a narrower distribution (Fig. 1b, c).

At the presynaptic terminal, Ca^2+^ entry through VGCCs activates kinases such as CaMKII, which phosphorylate a broad set of proteins critical for neurotransmitter release (24). These phosphorylation events regulate vesicle mobilisation, priming, and exocytosis, and a well-characterised example is synapsin, which releases synaptic vesicles from the reserve pool following CaMKII-dependent phosphorylation (25). To investigate if CaMKII activity is also required for calcium-dependent localisation of aSyn, we treated neurons with the CaMKII inhibitor KN-93 (26). Our results show that when CaMKII activity was inhibited, calcium no longer significantly altered aSyn-LTCC proximity (Fig. 1d, e), indicating that CaMKII activity is necessary for the calcium-driven positioning of aSyn.

While mean distances provide a measure of average molecular proximity, they do not capture changes in the shape of the distance distribution. Excess kurtosis was therefore calculated as a measure of tail heaviness relative to a normal distribution. Analysing the shape of spatial distributions of aSyn relative to LTCCs revealed substantial shifts in excess kurtosis across conditions (Table 1), with higher values indicating a sharper central peak together with rarer but more extreme tails. Specifically, an excess kurtosis of −0.51 was observed under the calcium-free condition, indicating a broad distribution of aSyn-LTCC distances and reduced central concentration. In contrast, both the spontaneous and stimulated conditions exhibited higher excess kurtosis values (9.56 and 8.7 respectively), reflecting a strong central concentration, driven primarily by an enrichment of very short aSyn-LTCC distances. This difference was less pronounced following the addition of KN-93, as excess kurtosis values for the calcium-free + KN-93 (3.01) and the spontaneous + KN-93 (4.84) conditions became more similar, suggesting reduced differences in the extent of close aSyn-LTCC associations under these buffer conditions. Notably, independent of KN-93 treatment, both stimulated conditions exhibited high excess kurtoses values (8.7 and 11.38 in the absence and presence of KN-93, respectively), indicating a persistent enrichment of closely associated aSyn-LTCC pairs under depolarising conditions. Together, these findings demonstrate that calcium influx and CaMKII activity are important for aSyn’s recruitment to LTCC-rich nanodomains.

**Table 1.**
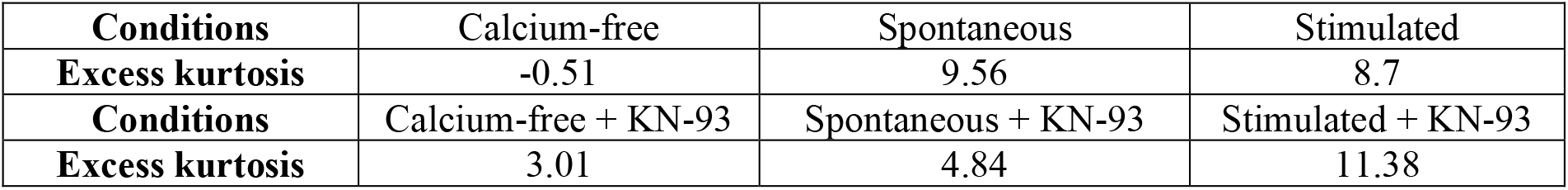
Excess kurtosis of spatial distributions of aSyn relative to LTCCs.

### Calcium increases presynaptic levels of total aSyn and pS129 aSyn and enhances C-terminal vesicle binding of pS129 aSyn

Phosphorylation of aSyn at serine 129 (S129) has previously been shown to increase in response to calcium influx, either through ionophores or VGCC activation (27,28). Several kinases can mediate this modification: casein kinases 1 and 2 phosphorylate aSyn efficiently in mammalian cells and in vitro (29), while CaMKII also leads to aSyn phosphorylation (30). Although S129 phosphorylation has been extensively studied in the context of aSyn aggregation, its potential role in regulating aSyn localisation and function at presynaptic terminals under physiological conditions remains unclear. We therefore asked whether calcium contributes to the enrichment of aSyn and pS129 aSyn at these sites. To address this, we selected synaptosomes, isolated, metabolically active presynaptic terminals for these experiments as they allow high-throughput imaging with minimal background from somata and debris (31). Synaptosomes purified from Sprague Dawley rat brains were incubated in buffers mimicking either calcium-free, spontaneous, or stimulated conditions and then imaged using DNA-PAINT to localise both total aSyn and pS129 aSyn (Fig. 2a). Quantitative PAINT analysis revealed significantly higher total aSyn and pS129 aSyn counts in synaptosomes incubated in buffers invoking spontaneous activity than under calcium-free conditions (Fig. 2b and c), indicating that both total and pS129 aSyn cluster at presynaptic sites in a calcium-dependent manner. Although not significant, under stimulated conditions, a lower level of pS129 aSyn but not total aSyn was observed, suggesting that a subset of phosphorylation-dependent presynaptic clustering of aSyn may be preferentially engaged under spontaneous activities.

**Figure 2.**
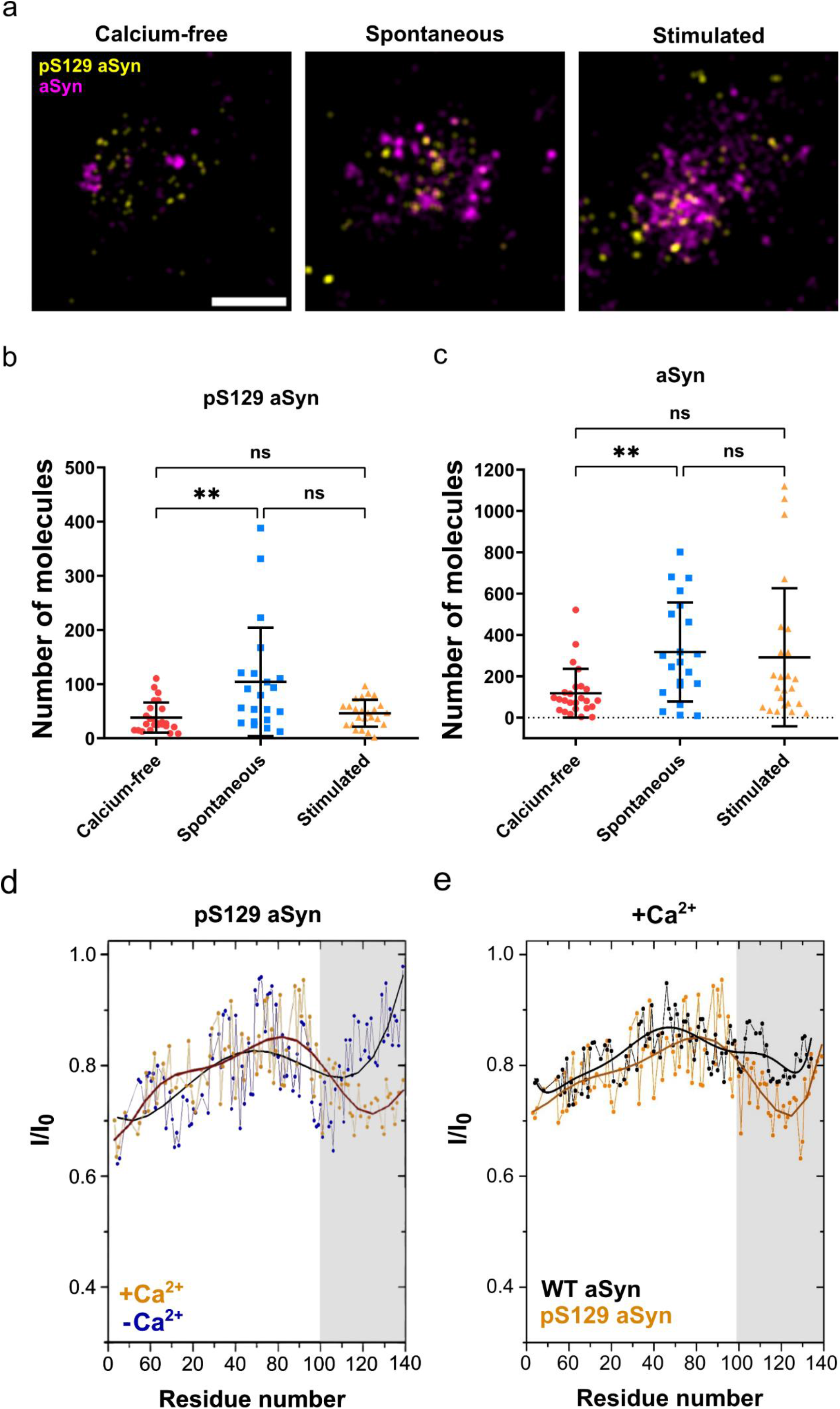
Calcium increases the abundance of both total and pS129 aSyn in synaptosomes and enhances the interaction of synaptic vesicles and pS129 aSyn at its C-terminus. **(a)** Representative reconstructed DNA-PAINT images of pS129 aSyn (yellow) and total aSyn (magenta) in synaptosomes under calcium-free (1 mM EGTA), spontaneous (2 mM CaCl_2_), or stimulated conditions (70 mM KCl and 2 mM CaCl_2_). Scale bar, 500 nm. **(b)** and **(c)**, qPAINT-quantified binding sites (molecule counts) for pS129 aSyn and total aSyn (mean ± s.d.) under three buffer conditions. Kruskal-Wallis test with Dunn’s multiple comparisons test. ns, not significant; **, P < 0.01. Data from N = 4 independent biological repeats were quantified. **(d)** CEST-NMR experiments were performed on pS129 aSyn in the presence of synaptic vesicles, both in the absence (blue) and presence (orange) of calcium (6 mM). **(e)** CEST-NMR experiments were performed on pS129 aSyn (orange) and wild-type (WT, black) aSyn (both 200 µM) in the presence of synaptic vesicles and calcium (6 mM).

We then investigated how calcium and S129 phosphorylation affect the interaction of aSyn and synaptic vesicles. In order to test this, we performed chemical exchange saturation transfer (CEST) in solution NMR using pS129 aSyn and synaptic vesicles isolated from rat brains in the presence or absence of calcium. CEST NMR measures how much the protein switches between free and vesicle-bound states (7), with a decrease in I/I_0_ indicating stronger interactions with synaptic vesicle. This analysis revealed that, in the absence of calcium, pS129 aSyn interacted most strongly with synaptic vesicles via the N-terminus. In the presence of calcium, however, the interaction at the C-terminus strongly increased (gray), while a slight increase was also observed at the N-terminus, as indicated by a decrease in I/I_0_ (Fig. 2d). Moreover, in the presence of calcium, pS129 aSyn showed stronger C-terminal interaction with synaptic vesicles compared to wild-type (WT) aSyn (Fig. 2e, gray). These findings suggest that aSyn interacts more strongly with synaptic vesicles at its C-terminus in the presence of calcium and S129 phosphorylation.

### aSyn and calcium support spontaneous DA release, and aSyn partitions to a small vesicle pool that are largely independent of canonical full-fusion associated recycling

There is ongoing debate regarding whether aSyn primarily regulates exocytosis or endocytosis, or more broadly modulates vesicle fusion behaviour at the presynapse in an activity-dependent manner (32). Under low levels of neuronal activity, synaptic vesicles can undergo kiss-and-run through transient flickering fusion, whereas strong or prolonged depolarisation typically increases the probability of full-fusion (33,34). Therefore, aSyn may differentially regulate specific vesicle populations depending on neuronal activity.

We first questioned whether aSyn and calcium were required for the release of DA under spontaneous and stimulated conditions. To test the contribution of calcium influx, we treated neurons with dihydropyridines (DHP), inhibitors of LTCCs that carry the majority of calcium conductance in substantia nigra (SN) dopaminergic neurons (35). Dopaminergic neurons were cultured from the midbrain of wild type (WT) and aSyn knock-out (SKO) mice and treated with DHP (5 μM isradipine) for 3 days to establish steady-state DA levels in vesicles. Neurons were then treated with 40 mM KCl for 5 min with or without 5 μM isradipine, followed by measurements of DA levels with high performance liquid chromatography with electrochemical detection (HPLC-EC). We found that DHP-treated cultures displayed a significant increase in intracellular DA levels in WT cells compared to no treatment (Fig. 3a). As LTCCs are involved in the generation of self-autonomous oscillatory activity of dopaminergic neurons (36–38), these results suggest that reduction or cessation of tonic firing leads to a buildup of stored neurotransmitter. Interestingly, no effect of DHP was observed in SKO cells, indicating that both aSyn and calcium influx are required for tonic (spontaneous) DA release. In contrast, no effect of DHP was found on DA release evoked by KCl in either WT or SKO neurons (Fig. 3b), in agreement with previous literature showing limited roles for LTCCs and aSyn in stimulated neurotransmission (22,39–41).

**Figure 3.**
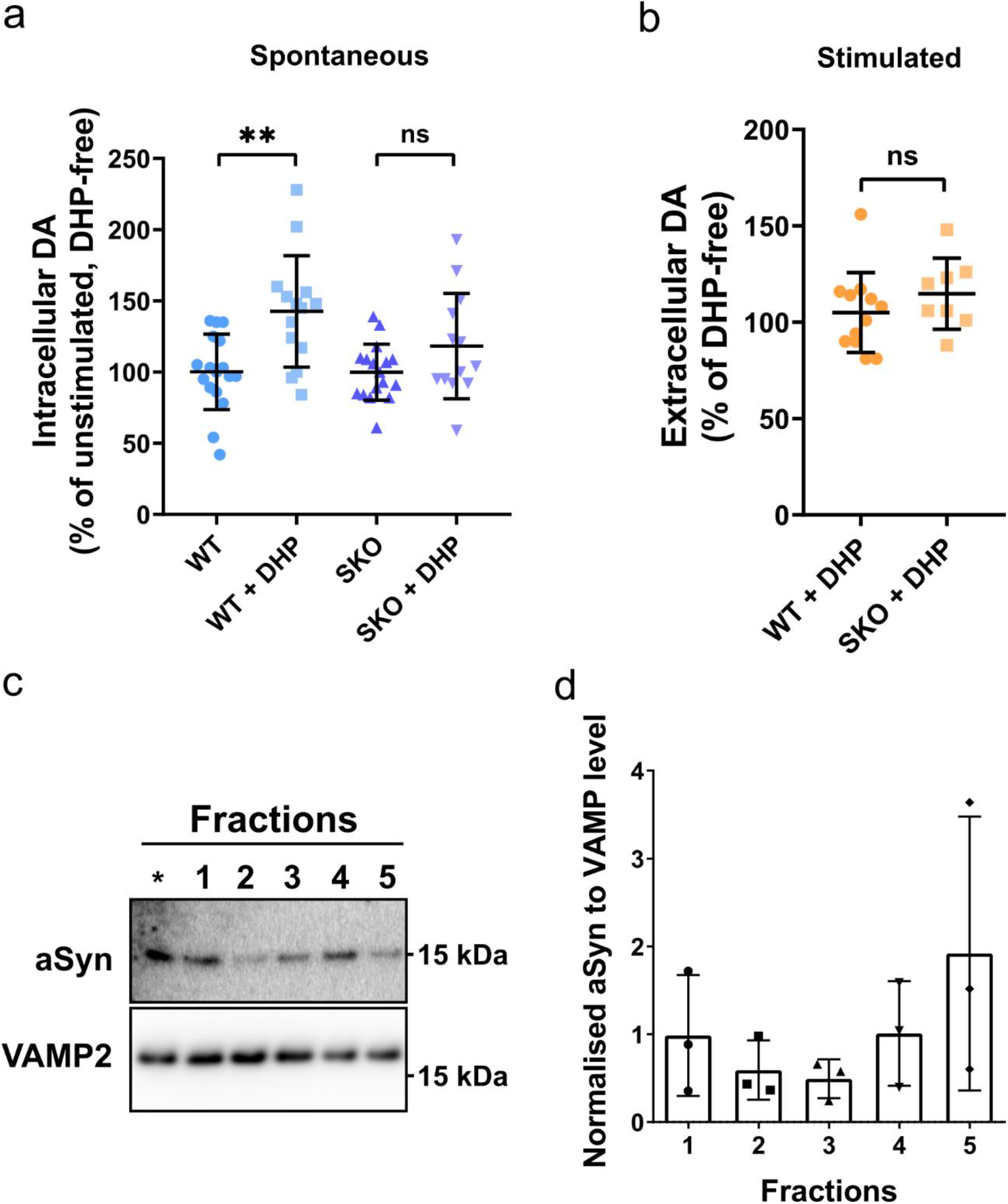
aSyn supports spontaneous LTCC-dependent DA release and associates with small synaptic vesicles. **(a)** Intracellular DA levels in WT and aSyn knock-out (SKO) neurons treated with dihydropyridines (DHP, 5 μM isradipine) for 3 days. Data shown as percentage change relative to genotype-matched controls. Brown-Forsythe ANOVA test with Tamhane’s T2 multiple comparisons test. ns, not significant; **, P< 0.01. Independent experiments/culture dishes (N/n) = 6/17 for WT, 5/14 for WT+DHP, 6/17 for SKO and 5/13 for SKO+DHP. **(b)** Effect of DHP treatment on DA release evoked by 40 mM KCl for 5 min. Extracellular DA levels in DHP-treated cultures were normalised to those in DHP-free sister cultures. Unpaired t test; ns, not significant. N/n = 4/12 for WT and 3/8 for SKO. **(c)** Representative Western blot images of aSyn and synaptobrevin 2 (VAMP2) from individual fractions of synaptic vesicles eluted from a size exclusion chromatography (SEC) column. aSyn more abundantly associated to vesicles in fractions eluted later from the column (higher-numbered fractions). A positive control (*) using brain homogenate showed presence of both proteins. **(d)** Quantification of the Western blots shows that aSyn is more abundant in vesicles eluting in later fractions. Data from N = 3 independent biological repeats were quantified.

Next, we asked whether aSyn associates with a distinct subpopulation of synaptic vesicles that are less engaged in canonical full-fusion. Given that aSyn has been shown to preferentially bind highly curved membranes (42), we hypothesised that it might be differentially enriched on smaller vesicles. To test this, we purified synaptic vesicles from WT Sprague Dawley rats and separated them by size-exclusion chromatography, where smaller vesicles elute later than larger ones. Western blot analysis of the fractions revealed that aSyn was enriched in vesicles eluting later from the column, whereas synaptobrevin 2 (VAMP2) was found in all fractions of vesicles eluting from the column (Fig. 3c and d). These results suggest that aSyn associates with a specific vesicle pool, likely corresponding to smaller vesicles.

To further test whether aSyn participates in vesicle pools that are less tightly coupled to full-fusion and canonical clathrin-mediated endocytosis (CME), we used direct dSTORM to analyse the spatial association of aSyn and VAMP2 with mCLING, a fluorescent tracer of endocytosis (Fig. 4a). Synaptosomes were incubated with mCLING in buffer that mimics the spontaneous neuronal activity at either 37 °C or 4 °C. At 37 °C, spontaneous exocytosis is allowed and endocytosed mCLING marks ongoing CME. In contrast, at 4 °C exocytosis is largely inhibited and CME is suppressed due to inhibition of clathrin uncoating, forcing membranes into a gel-like phase that raises the energetic barrier for endocytosis (43). After fixation, external mCLING was bleached with bromophenol blue to visualise only internalised tracers (44). Immunolabelling and dSTORM imaging for aSyn and VAMP2 was then performed (Fig. 4b), and a custom software was used to automatically detect and quantify single-molecule distributions in hundreds of synaptosomes.

**Figure 4.**
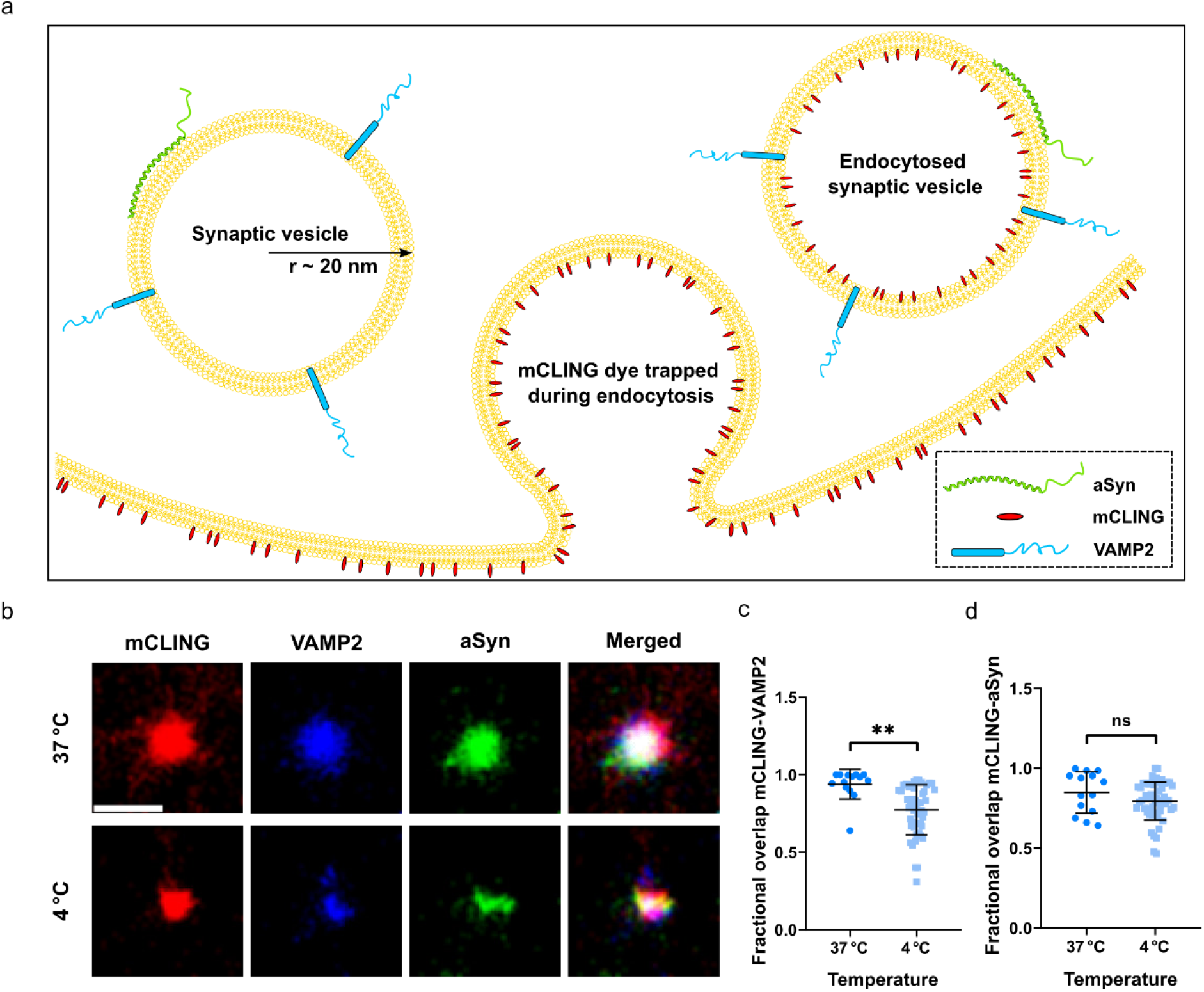
aSyn does not colocalise with the endocytic marker mCLING in a temperature dependent manner while VAMP2 colocalisation is temperature dependent. **(a)** Schematic of aSyn and VAMP2 on synaptic vesicles, and the mCLING staining after endocytosis. **(b)** Representative dSTORM images of synaptosomes stained with mCLING (red), VAMP2 (blue), aSyn (green), and an overlay of images. Scale bar, 500 nm. **(c)** and **(d)** Manders’ coefficients for mCLING colocalisation with VAMP2 or aSyn at 37 °C and 4 °C. Data from N ≥ 3 independent biological repeats were quantified. Kolmogorov-Smirnov test; ns, not significant; **, P < 0.01.

At 37 °C mCLING strongly colocalised with VAMP2 (Fig. 4c), often with Manders’ coefficients approaching 1, consistent with VAMP2 retrieval through CME to maintain v-SNARE recycling (45). In contrast, colocalisation between mCLING and aSyn was more heterogeneous (Fig. 4d), suggesting that aSyn is not tightly coupled to CME-dependent recycling. When experiments were repeated at 4 °C, mCLING-VAMP2 overlap decreased sharply (Fig. 4c), confirming the temperature sensitivity of CME, whereas mCLING-aSyn overlap was largely unchanged at 4 °C (Fig. 4d).

These observations support three conclusions: (i) aSyn is associated with vesicle pools that are not obligatorily recycled via CME; (ii) aSyn’s involvement in vesicle trafficking is not strongly temperature dependent, arguing against a role in mechanisms requiring large conformational rearrangements, such as CME; and (iii) aSyn function is not directly coupled to VAMP2, since the two proteins exhibit distinct colocalisation patterns with mCLING across conditions. Together, these findings indicate that aSyn acts on a specialised vesicle pool that is less tightly coupled to canonical full-fusion and CME-dependent recycling.

## Conclusion

Using DNA-PAINT super-resolution imaging, we found that calcium promotes aSyn clustering around LTCCs, with CaMKII contributing to this recruitment under spontaneous activity. In parallel, calcium increases endogenous total and pS129 aSyn at synaptosomes under spontaneous conditions. Additionally, NMR experiments showed that both calcium and S129 phosphorylation enhance aSyn’s affinity for synaptic vesicles. Functional assays further revealed that both LTCC-dependent calcium entry and aSyn appear to be necessary for DA release during spontaneous neuronal activity, but not under stimulation. Finally, biochemical fractionation and multi-colour single-molecule imaging showed that aSyn preferentially associates with small vesicles that are largely independent of full-fusion recycling pools.

Together, our findings support a model in which aSyn regulates presynaptic vesicle organisation and DA release through a calcium- and phosphorylation-dependent mechanism that largely operates outside of canonical exocytotic and recycling pathways. Rather than acting as a core component of full-fusion mediated neurotransmission, aSyn appears to fine-tune presynaptic function under conditions where calcium fluctuations are modest and canonical mechanisms are less engaged.

The increased enrichment of aSyn at LTCC microdomains under both spontaneous and stimulated conditions highlights its intimate coupling to calcium entry, consistent with the extended double-anchor model in which the N-terminus binds vesicle membranes while the C-terminus engages vesicles in a calcium-dependent manner (10). This arrangement may enable aSyn to tether vesicles at active zones, cluster them near calcium channels, and remodel local curvature, thereby facilitating a calcium-tuned release mechanism. Additionally, CaMKII activity, which contributes to aSyn phosphorylation (30), is required for aSyn clustering around LTCCs under spontaneous conditions, while pS129 species showing enhanced association with vesicles at both termini in the presence of calcium. This regulation suggests that S129 phosphorylation acts as a molecular switch, enabling aSyn to engage vesicle pools under defined activity states.

Importantly, this role is not apparent during strong stimulation, when full-fusion exocytosis predominates. Blockade of LTCC reduces DA release in WT but not aSyn knockout neurons under spontaneous but not strongly stimulated conditions, indicating that LTCC-dependent DA release requires aSyn. Moreover, LTCC activity is known to be engaged during autonomous pacemaking in midbrain dopaminergic neurons (19), further emphasising the relevance of this pathway under basal activity states. Together, these observations point to aSyn’s involvement in spontaneous or restricted fusion modes, such as kiss-and-run through transient flickering fusion, that are potentially overshadowed under conditions of robust neuronal firing or in cells with overexpression of aSyn.

Although aSyn has been proposed to modulate SNARE function through interactions with VAMP2 (46), our data suggest an additional mechanism. aSyn abundance was increased on smaller vesicles, while VAMP2 remained uniformly distributed, suggesting that aSyn preferentially associates with a subset of synaptic vesicles. Furthermore, aSyn did not exhibit the temperature-dependent endocytic behaviour characteristic of VAMP2, despite remaining colocalised on vesicles. This dissociation suggests that aSyn is not obligately coupled to canonical SNARE/CME-associated recycling pathways. Instead, aSyn may exploit its intrinsic ability to induce membrane curvature (47), thereby modulating vesicle fusion behaviour through alternative mechanisms under conditions where dominant full-fusion pathways are less effective or saturated. Such conditions may favour flickering fusion events described in dopaminergic neurons (48), when synaptic vesicles form and maintain a highly curved Ω-shaped membrane profile (49). This view of aSyn as a conditional regulator of presynaptic vesicle pools is further supported by its regulation in vivo, potentially contributing to synaptic plasticity (50). aSyn expression is upregulated during song learning in birds (51), a paradigm of sustained neuronal activity in which conventional release mechanisms may be insufficient to meet demand.

These insights have important implications for Parkinson’s disease. While S129 phosphorylation of aSyn has long been associated with pathology, our findings indicate that pS129 may also reflects physiological regulation of vesicle dynamics. Thus, therapeutic approaches aiming to reduce aSyn levels or block its phosphorylation must account for potential disruption of its normal presynaptic functions. More broadly, defining aSyn as a calcium- and phosphorylation-tuned modulator of vesicle organisation and DA release provides a conceptual framework for reconciling its physiological role with its pathological aggregation in disease. In this light, PD may involve not only toxic gain-of-function effects of aggregated aSyn, but also loss of its finely tuned contributions to presynaptic vesicle regulation.

In summary, our findings identify aSyn as a calcium- and phosphorylation-tuned regulator of presynaptic vesicle organisation and DA release. Rather than acting as a core component of full-fusion, aSyn modulates spontaneous DA release by promoting flickering fusion events by clustering small vesicles near LTCCs through a double-anchor mechanism. This function is masked during strong stimulation, when canonical pathways dominate, but may be essential under conditions of sustained activity or limited vesicle supply. Importantly, phosphorylation at S129, often regarded solely as a pathological marker, also reflects physiological regulation, highlighting the need to consider potential loss of normal aSyn function when developing therapeutic strategies for Parkinson’s disease.

## Materials and Methods

### Synaptosome isolation and treatment for DNA-PAINT

The animal procedure was approved by the University of Cambridge, University Biomedical Services, AWEBR Committee. Synaptosomes were purified from Sprague-Dawley rats (Charles River) with a Percoll gradient procedure by using different centrifugation steps as previously described (52). Briefly, all preparations were done on ice or at 4°C unless stated, the brain was homogenised in a glass-Teflon EUROSTAR20 homogeniser (IKA, Oxon, UK) in homogenising buffer made from sucrose/EDTA buffer (320 mM sucrose, 1 mM EDTA, 5mM Tris, pH 7.4) with 50 mM DTT, using 10 strokes at 800 rpm. Synaptosomes were isolated using 3-23% Percoll gradients. The synaptosome containing fractions were pooled and resuspended in extracellular buffer solution (130 mM NaCl, 5 mM KCl, 20 mM 4-(2-hydroxyethyl) piperazine-1-ethanesulfonic acid (HEPES) sodium salt, 5 mM NaHCO_3_, 1.2 mM Na_2_HPO_4_, 1 mM MgCl_2_ and 10 mM Glucose). µ-slide 8 well high glass-bottom plates (ibidi GmbH) were cleaned with 1 M KOH for 30 min, washed with Tris-buffered saline (TBS) twice and coated with 0.1% (w/v) poly-L-lysine (PLL) before the adhesion of synaptosomes. After incubating on ice for 10 min, unadhered synaptosomes were removed. Adhered synaptosomes were then treated with 200 µL of three different buffer solutions at 37℃ for 30 min: extracellular buffer solutions added by (1) 1 mM ethylene glycol-bis (β-aminoethyl ether)-N, N, N’, N’-tetraacetic acid (EGTA) (calcium-free condition), (2) 2 mM CaCl_2_ (spontaneous condition), or (3) 2 mM CaCl_2_ and 70 mM KCl (stimulated condition). After treatment, synaptosomes were fixed with 4% formaldehyde and 1% glutaraldehyde in TBS for 10 min at room temperature. To quench fixation, 0.1 M glycine in TBS was added for 5 min followed by washing with TBS once.

### Fluorescent labelling of aSyn, VAMP2 and mCLING in synaptosomes

Following isolation and plating in µ-slide 8 well high glass-bottom plates, adhered synaptosomes were incubated for 30 min at 37 °C or 4°C with extracellular buffer solutions containing 0.5 nmol/mL mCLING-ATTO647N (#710 006AT647N, SYnaptic SYstems, Göttingen, Germany), and 1 mM EGTA (calcium-free condition) or 2 mM CaCl_2_ (spontaneous condition). Synaptosomes were then fixed with 4% formaldehyde and 0.2% glutaraldehyde (Sigma-Aldrich) in PBS. Fixation was quenched by washing with 0.1 M glycine in PBS for 5 min. To chemically quench the mCLING that bound to the outer membrane of the plasma membrane of the synaptosomes, 0.75 mM bromophenol blue was incubated for 5 min and washed off with PBS (44). Fixed synaptosomes were permeabilised in 0.1% Triton X-100 in PBS and blocked for 1 h in 5% goat serum in PBS. Immunostaining was performed using primary antibodies against aSyn (1:500, D37A6, Cell Signalling Technology) and VAMP2 (1:500, 104211, SYSY) in blocking buffer overnight at 4 °C. After washing in PBS, secondary antibodies, anti-rabbit Alexa Fluor 568 (1:500, ab150067, Abcam) and anti-mouse Alexa Fluor 488 (1:500, ab175700, Abcam) were applied for 1 h at room temperature. Labelled samples were washed, mounted on #1.5H precision coverslips, and imaged in photoswitching buffer during multi-colour dSTORM acquisition.

### Rat primary dopaminergic neuronal cell cultures and treatment

Primary cortex glia cells were isolated from Sprague-Dawley rats (Charles River) as previously described (53). 130,000 cells were plated with glia plating media (90% Dulbecco’s Modified Eagle’s Medium-high glucose (DMEM), 10% fetal bovine serum (FBS), 2 mM GlutaMAX supplement, and 1X antibiotic-antimycotic) in each well of the µ-slide 8 well high glass-bottom plates (ibidi GmbH) coated with 0.01% (w/v) PLL. One day after plating, unattached cells were removed by vigorous pipetting, and the media was changed twice a week until cells were confluent. Cell division was stopped by changing to glia stopping media (90% DMEM, 5% FBS, 2 mM GlutaMAX supplement, 1X antibiotic-antimycotic, and 100 μM cytarabine). On the next media changing day, microglia were removed with vigorous pipetting, and glia maintaining media (glia stopping media without cytarabine) was used for the following media change, until cells are confluent and ready for plating dopaminergic neurons. The cell incubator was set at 37°C with a 5% CO_2_ atmosphere.

Primary midbrain dopaminergic neurons were isolated from postnatal day 2 (P2) Sprague-Dawley rats (Charles River) and purified with OptiPrep Density Gradient Medium (Sigma-Aldrich) before being plated on glia feeder layers as previously described (53). 7,500 cells were plated with postnatal neuron media (Neurobasal A medium supplemented with 1X B-27, 0.5 mM GlutaMAX supplement, and 10 μg/mL gentamicin) in each well and 10 ng/mL GDNF (glial cell-derived neurotrophic factor) was added as a growth factor. The day after plating, 10 μM cytarabine was added to avoid glia growth. Half the media was changed once a week until 14 days in vitro (DIV).

At 14 DIV, cells were treated with 300 µL of three different buffer solutions: extracellular buffer solutions added by 1 mM EGTA, 2 mM CaCl_2_, or 2 mM CaCl_2_ and 70 mM KCl at 37℃ for 30 min, in the presence or absence of 20 μM KN-93 (water soluble, ab120980, Abcam), an inhibitor of calcium/calmodulin-dependent kinase type II (CaMKII). After treatment, cells were fixed with 4% formaldehyde in phosphate-buffered saline (PBS) at 37℃ for 10 min followed by washing 3 times with PBS.

### Synaptosomes and dopaminergic neurons immunolabeling for DNA-PAINT

Fixed samples were permeabilised with 0.05% (v/v) PBS-Tween 20 (PBST) for 5 min and then blocked with 5% donkey serum (Sigma-Aldrich) in PBST for 1 hour at room temperature. Synaptosomes were then incubated with mouse anti-pS129 aSyn (1:200, lab-made, BL pS129M) and rabbit anti-aSyn (1:200, Synaptic Systems, 128002), while dopaminergic neurons were labelled with mouse anti-aSyn (1:500, Thermo Fisher Scientific, LB509) and rabbit anti-Ca_V_1.3 (CACNA1D) (1:200, Alomone Labs, ACC005) primary antibodies for 1 hour at room temperature. After washing with PBS 3 times, samples were incubated with 2.5 μg/mL DNA-conjugated polyclonal donkey anti-mouse (Massive Photonics) and DNA-conjugated polyclonal donkey anti-rabbit (Massive Photonics) secondary antibodies for 1 hour at room temperature. After washing with PBS 3 times, samples were kept in PBS with 0.1% NaN_3_ at 4°C.

### Mouse midbrain neuron cell cultures, DHP treatment, and HPLC-EC

The use of animals followed the National Institutes of Health guidelines and was approved by the Institutional Animal Care and Use Committee of Columbia University and New York State Psychiatric Institute. Mice were housed under a 12/12h light/dark cycle with *ad libitum* access to food and water. aSyn null mice and their isogenic mixed background controls (RRID: IMSR_JAX:003692 and B6129SF2/J, IMSR_JAX:101045) were from JAX.

Midbrain dopaminergic neurons from postnatal day 0-2 mice of both sexes were dissected, dissociated, and plated on a monolayer of rat cortical astrocytes at the plating density of ~100,000 cells/cm^2^, as described previously (54). Isradipine (5 μM, Tocris) was added 12-15 days after plating daily for 3 days. On the day of the experiment, cells were rinsed with PBS, which was then changed to Tyrode’s saline containing vehicle, 5 μM isradipine, 40 mM KCl or both isradipine and KCl. After 5 min incubation at 37°C, supernatant was collected and immediately mixed with 0.2 M perchloric acid (1:1 volume) to deproteinise the sample and prevent dopamine auto-oxidation. Perchloric acid was also added into the wells with cells to measure intracellular DA concentration. After 10 min incubation at room temperature, samples were centrifuged at 10,000 g for 5 min at 4°C. Supernatant was collected, stored at −80°C and analysed with high performance liquid chromatography with electrochemical detection (HPLC-EC) within the following two weeks. DA concentrations in each group of samples were normalised to the levels in the corresponding control group.

### Recombinant overexpression and purification of ^13^C and ^15^N labelled human WT aSyn

The recombinant overexpression and purification of double-labelled human WT aSyn were carried out as described (55). pT7 plasmid encoding hWT aSyn was used for transformation in BL21(DE3) *E. coli* cells on an ampicillin agar plate. A single colony was transferred to 200 mL of Luria broth (LB) medium containing ampicillin (100 μg/mL; small-scale culture; AppliChem, A0839) and incubated overnight at 37 °C and 180 rpm. The following morning, the LB-grown bacterial cells were pelleted. These cells were then transferred to minimal medium containing ^13^C-labelled glucose and ^15^N-labelled ammonium chloride, with a starting OD600 of 0.1, for the production of ^13^C, ^15^N-labelled WT protein before induction. After induction using 1 mM 1-thio-β-d-galactopyranoside (AppliChem, A1008) upon cells reaching OD600 at 0.6 and following another 5 hours incubation at 37 °C and 180 rpm, bacteria were lysed by sonication, and the lysate was subjected to 100°C boiling for ~15 min and further followed by the next steps of purification protocol such as anion exchange chromatography, reversed-phase HPLC, and lyophilization as described previously (55). The purity of the lyophilised ^13^C, ^15^N-labelled human WT aSyn was analysed by ultraperformance liquid chromatography (UPLC) and electrospray ionization mass spectrometry (ESI-MS) (Supplementary Fig 2).

### Preparation of pS129-phosphorylated ^13^C/^15^N human aSyn by enzymatic phosphorylation

Purified, lyophilised, double-labelled (^13^C/^15^N) WT aSyn was prepared as previously described (56). Briefly, 500 µg of protein was weighed using an analytical microbalance and dissolved in 195 µL of freshly prepared phosphorylation buffer (50 mM HEPES, 1 mM MgCl_2_, 1 mM EGTA, 1 mM DTT). Phosphorylation was initiated by the addition of 4 µL of 100 mM Mg-ATP (final concentration 2 mM) and 0.42 µg PLK3 (1 µL). The reaction mixture was gently mixed by pipetting (without vortexing) and incubated for 12 h at 30 °C without agitation. Phosphorylation efficiency was assessed by mass spectrometry.

### LC-MS (ESI-LTQ) and UPLC characterisation of S129-phosphorylated ^13^C/^15^N human aSyn

For protein characterisation, liquid chromatography-mass spectrometry (LC-MS) using an LTQ instrument (Thermo Scientific) with electrospray ionisation was used (55,57). Samples were run using a Poroshell 300SB-C3 column (1.0 × 75 mm, 5 µm; Agilent Technologies) via a linear gradient from 5% to 95% acetonitrile (0.1% formic acid) over 10 min at a flow rate of 300 µL/min. Mass spectra were processed using MagTran software (Supplementary Fig 3). UPLC analysis was performed on a Waters Acquity H-Class system using a C4 column (300 Å, 1.7 µm, 2.1 × 150 mm) with UV detection at 214 nm. Samples were separated using a gradient from 10% to 90% acetonitrile at a flow rate of 0.6 mL/min over 3 min (Supplementary Fig 3).

### Synaptic vesicle isolation

Isolation of synaptic vesicles (SVs) from two rat brains per preparation was performed as previously described (58). Brains from two euthanized Sprague-Dawley rats were removed and washed in ice cold homogenising buffer (320 mM sucrose, 4 mM HEPES, EDTA free cOmplete protease inhibitor, pH 7.4), before being homogenised in 9 mL homogenising buffer using a glass-Teflon EUROSTAR20 homogenizer (IKA, Oxon, UK) for 10 strokes at 900 rpm. The mixture was centrifuged at 1000 x g for 10 min at 4°C. The supernatant was collected and centrifuged at 15, 000 x g for 15 min at 4°C and stored on ice, while the pellet was resuspended in 1 mL homogenising buffer. 9 mL of ice-cold ddH_2_O with EDTA free cOmplete protease inhibitors was added to the resuspended pellet and homogenised for three strokes at 2000 rpm. 50 µL of 1 M HEPES(NaOH) was immediately added after homogenising. The homogenate was centrifuged at 17, 000 x g for 15 min at 4°C. The resulting supernatant was combined with the supernatant on ice. The combined supernatants were centrifuged at 48, 000 x g for 25 min at 4°C. The supernatant was collected and homogenised for five strokes at 2000 rpm and drawn up and dispersed through a 30-gauge needle to disperse vesicle clusters. The supernatant containing vesicles was divided and 5 mL layered over a 5 mL 0.7 M sucrose cushion. The sucrose cushions were centrifuged at 133,000 x g for 1 hour at 4°C. 500 µL fractions from the cushions were removed starting at the top of the gradient. 5 µL of each fraction was spotted onto a PVDF membrane and used for a dot blot. Fractions 12-20 were pooled and centrifuged at 300,000 x g for 2 hours. The vesicle pellet was resuspended in 1 mL column buffer (100 mM Tris-HCl pH 7.6, 100 mM KCl) and homogenised in a 1 mL glass-teflon homogeniser for 10 strokes at 900 rpm. The vesicles were drawn up through and expelled through a 30-gauge needle three times. The vesicles were loaded onto a pre-prepared Sephacryl S-1000 column and a peristaltic pump with a flow rate of ~6 mLh^-1^ was set up in a cold room, 4°C. Fractions of 0.7 mL were collected and analysed on a nanodrop to determine protein concentration.

### Western blot analysis of vesicle fractions

Synaptic vesicles were separated by size using a size exclusion column (SEC). Fractions were collected, and protein content was monitored by absorbance at 280 nm (A280). Synaptic vesicles eluted in the second peak on the chromatograph. Each fraction from this peak was centrifuged, and the supernatant was removed to leave approximately 10 uL. The fractions were then snap-frozen in liquid nitrogen and lyophilised. Fractions were incubated with 20 uL of LDS buffer and analysed by western blot following transfer onto PVDF membranes. The membrane was first probed with anti-aSyn primary antibody (Cell Signalling Technology, D37A6, 1:1000) and HRP-conjugated anti-rabbit IgG secondary antibody (GE Healthcare, 1:4000). Blots were subsequently stripped and reprobed with anti-synaptobrevin (VAMP2) primary antibody (SYSY, 104211, 1:1000) followed by an HRP-conjugated anti-mouse IgG secondary antibody (GE Healthcare, 1:4000). (The raw Western blot images from 3 independent experiments are provided in Supplementary Fig 1) Densitometric analysis of the western blots was used to determine the percentage of aSyn present in vesicle-enriched fractions, normalised to VAMP2, which is abundantly found on synaptic vesicles (59).

### Multi-colour dSTORM imaging of synaptic proteins in synaptosomes

Single-molecule localisation microscopy (direct STORM, dSTORM) was performed on a custom-built wide-field microscope based on an Olympus IX-73 frame (Olympus, Center Valley, PA). Excitation was provided by three lasers: 488 nm (Coherent Sapphire 488-300 CW CDRH), 561 nm (Cobolt Jive 500 561 nm), and 647 nm (MPB Communications VFL-P-300-647-OEM1-B1). Laser beams were reflected from a multi-band dichroic mirror (Chroma ZT488/561/647rpc) and focused through a 100× 1.49 NA oil-immersion objective (Olympus UAPON100XOTIRF). Fluorescence emission passed through the dichroic and a set of 25 mm band-pass filters (Semrock FF01-525/45-25, Semrock FF01-600/37-25, and Semrock FF01-680/42-25 respectively for the 488, 561, and 647-nm laser lines) before being relayed to a camera (Andor iXon Ultra 897) via a Cairn Twincam image splitter (1.3× magnification). The reflective arm of the splitter was unused during acquisition. To obtain uniform excitation across the field of view, the system employed a Gaussian-to-top-hat beam shaper (Topag Lasertechnik GTH-5-250-4-VIS) (60). Highly inclined laminated optical sheet (HILO) illumination was implemented by laterally offsetting the excitation beam in the objective’s back focal plane (61). Sequential imaging was performed for the 647 nm, 561 nm, and 488 nm channels to minimise spectral crosstalk and photobleaching. For each channel, 16,000 frames were recorded at 10 ms exposure per frame, with an electron-multiplying (EM)-gain of 200.

### dSTORM image reconstruction and data analysis

Acquired image stacks were reconstructed using ThunderSTORM plugin in ImageJ (62). To automate and standardise quantitative analysis, a custom MATLAB-based toolkit named SynaptoAnalysis was developed. The software processes localisation tables exported from ThunderSTORM to extract spatial statistics describing the organisation of synaptic proteins at the nanoscale. The workflow consisted of four main stages: (1) chromatic registration to correct for spectral misalignment; (2) automatic detection of individual synaptosomes; (3) quantitative multicolour colocalisation analysis; and (4) measurement of protein clustering. All source code and documentation are publicly available at https://github.com/pedropabloVR/Synaptosome-Analysis.

#### Chromatic registration

Chromatic aberrations between emission channels were corrected using 0.1 μm multicolour fluorescent beads (TetraSpeck Microspheres, Invitroge). Localisations were first extracted in ThunderSTORM and then aligned in MATLAB by mapping coordinates from the 488 nm and 561 nm channels to a 647 nm reference using a locally weighted polynomial transformation (MATLAB fitgeotrans function). The registration accuracy was evaluated using the total registration error (TRE), defined as the mean Euclidean distance between fiducial centroids before and after correction. TRE was reduced from ~35-57 nm pre-correction to ~8-13 nm post-correction, consistent with accepted chromatic alignment performance for multicolour single-molecule localisation microscopy (SMLM).

#### Automatic detection of synaptosomes

Localisation tables were filtered to remove low-quality detections based on photon count, Gaussian fit sigma, and localisation precision (thresholds defined empirically for each dataset). Noise and background were further suppressed using a nearest-neighbour density filter and Otsu’s adaptive thresholding. Candidate synaptosomes were segmented based on overlapping fluorescence from the mCLING membrane dye and synaptic protein labels (aSyn and VAMP2). Objects showing less than 10% area overlap or fewer than 200 localisations were discarded. To prevent redundant detection, candidates located within 300 nm of each other were excluded. Validated regions (80 × 80 pixels^2^) were cropped and exported as individual localisation files for further analysis.

#### Colocalisation analysis

Spatial colocalisation between proteins was quantified using Manders’ overlap coefficients (M_1_, M_2_), which report the fraction of fluorescence from one channel overlapping with another. A Weighted Overlap Coefficient (WOC) was also computed as a combined metric of spatial co-occurrence. These analyses were implemented in MATLAB and applied to compare aSyn and VAMP2 distributions relative to internalised vesicles labelled with mCLING under varying calcium and temperature conditions.

#### Cluster size measurement

Protein clustering within each synaptosome was quantified using the root-mean-square distance from centroid (RMSD) metric, defined as the square root of the mean squared radial distance of localisations from their cluster centroid. This measure provides an estimate of the spatial spread of localisations within a cluster and was benchmarked against Ripley’s K-function and its normalised form, Ripley’s H-function (63). Synthetic SMLM datasets generated from a computational model of a synaptosome were used to validate the RMSD method, confirming that it is less sensitive to variations in localisation density and faster to compute than Ripley’s statistics, making it well suited for high-throughput SMLM analysis.

#### Implementation and computational performance

All algorithms were implemented in MATLAB R2019a (MathWorks Inc., Natick, MA, USA). Each SMLM dataset contained approximately 1-2 million localisations. Typical processing times ranged between 30 and 90 min per dataset, depending on synaptosome density. The density-filtering and neighbour-search steps were the most computationally demanding elements of the pipeline.

### DNA points accumulation for imaging in nanoscale topography (DNA-PAINT)

Before imaging, PBS in each well was replaced with imager strands (Massive Photonics) diluted in 100 μL imaging buffers (Massive Photonics). For synaptosomes, 1 nM imager strands conjugated to ATTO565 or ATTO655 were used, while for dopaminergic neurons, 2.5 nM imager strands conjugated to Cy3B, and 0.25 nM imager strands conjugated to ATTO655 were used (all from Massive Photonics). DNA points accumulation for imaging in nanoscale topography (DNA-PAINT) imaging was performed on an inverted Olympus IX-73 microscope (Olympus Corporation) equipped with a TIRF oil-immersion objective (Apochromat 100×, NA 1.49 Oil, Olympus). Excitation was provided by either a 561 nm or 647 nm laser, combined using a dichroic mirror (ZT647rpc, Chroma) and directed onto the sample with a HILO illumination. Emitted fluorescence was collected in transmission through a dichroic filter cube (Cairn OptoSpin) in transmission and a wavelength-specific emission filter (BrightLine, 600/37 or 680/42) and detected using an electron-multiplying charge-coupled device (EMCCD) camera (Andor iXon Ultra 897). A total of 15,000 image frames were recorded at different exposure times (0.1 s or 0.2 s for synaptosomes, and 0.05 s for dopaminergic neurons) within a field of view (FOV) of 256 × 256 camera pixels. Before switching imager strands, samples were washed with 10 mL PBST to remove the previous imager strand. For drift correction, 0.1 μm multicolour fluorescent beads were imaged in the 512 × 512 FOV both before and after each wash step.

### DNA-PAINT image reconstruction and analysis

To account for dichroic effects of the microscope and to register the optical channels, images of 0.1 μm multicolour fluorescent beads were acquired in all channels. Bead positions were extracted using the ThunderSTORM plugin in ImageJ. Using these positions, transformation matrices between the reference channel and the others were computed. Single-molecule localisations were extracted from each of the 15,000-frame images using the ThunderSTORM plugin in ImageJ and subsequently registered across channels using the computed transformation matrices. For multicolour DNA-PAINT, changing the imaging medium and washing the sample between rounds introduced additional drift. To correct for this drift, sparsely distributed multicolour fluorescent beads were placed to ensure at least a few were present in each field of view. Images of these beads were acquired immediately before and after washing, and the drift was corrected. Following reconstruction and registration, individual synaptosomes were identified, isolated, cropped, and plotted using the custom MATLAB-based toolkit, SynaptoAnalysis.

#### qPAINT

Synaptosome regions (clustered localisations) were cropped to 150 × 150 pixels, and the number of binding sites for total aSyn and pS129 aSyn were quantified using a custom-written MATLAB script based on the qPAINT method (64). In brief, qPAINT allows to estimate number of binding sites in a given region from known experimental parameters, such as imager strand concentration (c), exposure time and association rate of strands (*K*_on_). The cumulative intensity function was fitted to an exponential decay function, from which the mean dark time (*τ*_*d*_) was determined. The number of binding sites was subsequently calculated as 1/(*τ*_*d*_ *c K*_on_).

#### Nearest neighbour distance

To assess colocalisation between aSyn and LTCC in dopaminergic neurons, the nearest neighbour distance was calculated using a custom-developed MATLAB application, ColocAnalyser, that was developed in our lab to measure a number of different colocalisation metrics (https://github.com/StasMakarchuk/ColocAnalyzer). To be brief, reconstructed DNA-PAINT images were pre-processed using median filtering and Otsu’s thresholding. For each localisation in the aSyn channel, the distance to the closest spot in the LTCC channel was measured. Distances ranging from 0 to 2000 nm were binned into histograms with a bin width of 100 nm. To capture the variation in data distribution between the conditions without data transformation, excess kurtosis was derived using a basic stats calculator (65).

### Chemical exchange saturation transfer (CEST) NMR

Chemical exchange saturation transfer (CEST) solution nuclear magnetic resonance (NMR) experiments were employed to directly probe the equilibrium between vesicle-unbound and vesicle-bound states of aSyn. Compared to standard heteronuclear correlation spectroscopy, CEST provides enhanced sensitivity for characterizing the equilibrium between NMR-visible (unbound aSyn) and NMR-invisible (vesicle-bound aSyn) species, with significant sensitivity even at low vesicle-to-protein ratios. In this approach, saturation of the bound state is transferred to the free state through conformational exchange between the two species, leading to attenuation of the peak intensities corresponding to the NMR-visible unbound form. CEST experiments were carried out at 10 °C on a Bruker spectrometer operating at ^1^H frequencies of 700 MHz equipped with triple resonance HCN cryo-probe. The measurements were based on ^1^H-^15^N HSQC experiments by applying constant wave saturation of 400 Hz in the ^15^N channel. Saturations were applied at −1.5 and +1.5 kHz, whereas an additional spectrum, saturated at −100 kHz, was recorded as a reference. The CEST experiments were recorded using a data matrix consisting of 2048 (*t*_2_, ^1^H) × 220 (*t*_1_, ^15^N) complex points.

### Statistical analysis

Statistical analyses were performed using GraphPad Prism 8. Data are presented as mean ± standard deviation (SD). Normality of data distribution was assessed using the D’Agostino and Pearson Test or the Shapiro-Wilk test (for small sample sizes). Depending on data characteristics and experimental design, statistical significance was evaluated using Brown-Forsythe and Welch ANOVA or Kruskal-Wallis tests for multiple-group comparisons, and unpaired t tests or Kolmogorov-Smirnov tests for pairwise comparisons. Significance levels are indicated as follows: ns, not significant; *, P < 0.05; **, P < 0.01. Quantification was based on data from at least N = 3 independent experiments. Specific statistical tests used for individual experiments are detailed in the corresponding figure legends.

## Acknowledgements

G.S.K.S. acknowledges funding from the Wellcome Trust (065807/Z/01/Z) (203249/Z/16/Z), the UK Medical Research Council (MRC) (MR/K02292X/1), Alzheimer Research UK (ARUK) (ARUK-PG013-14), Michael J Fox Foundation (16238 and 022159), and Infinitus China Ltd. C.F.K. acknowledges funding from the UK Engineering and Physical Sciences Research Council (EP/L015889/1 and EP/H018301/1), the Wellcome Trust (3-3249/Z/16/Z and 089703/Z/09/Z), the UK Medical Research Council (MR/K015850/1 and MR/K02292X/1), and Infinitus China Ltd. N.F.L. acknowledges the Swiss National Science Foundation (Grant Number P2EZP2 199843).

