## Supplementary Figures for "α-Synuclein Facilitates Spontaneous Dopamine Release in a Calcium- and Phosphorylation-Dependent Manner"

### Supplementary information

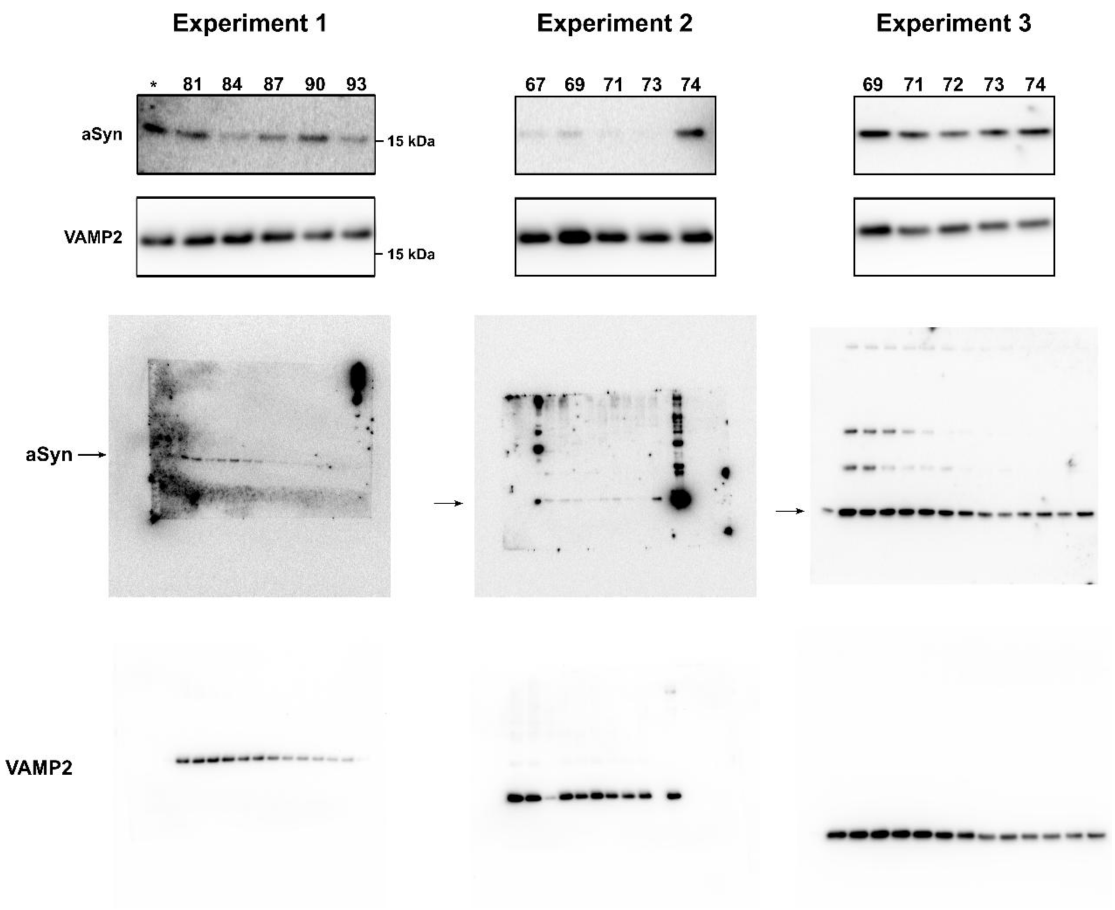


**Supplementary Fig 1 | Original Western blot images from 3 independent experiments corresponding to data shown in Fig. 3c and d.**

Mass spectrometry

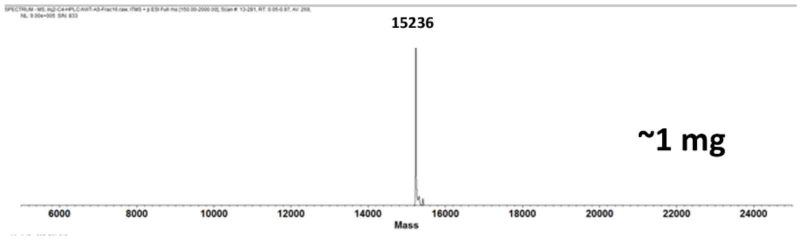


UPLC

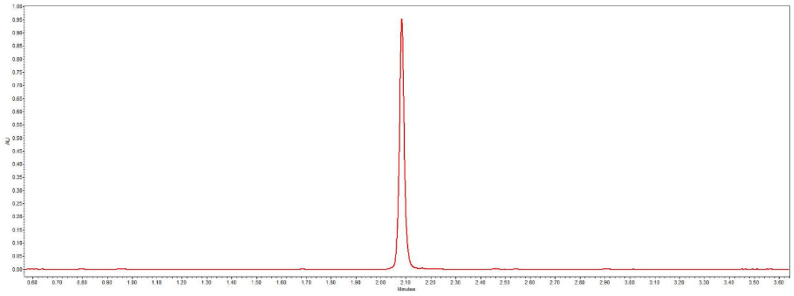

**Supplementary Fig 2 | ESI-MS and UPLC characterization of WT human aSyn.**

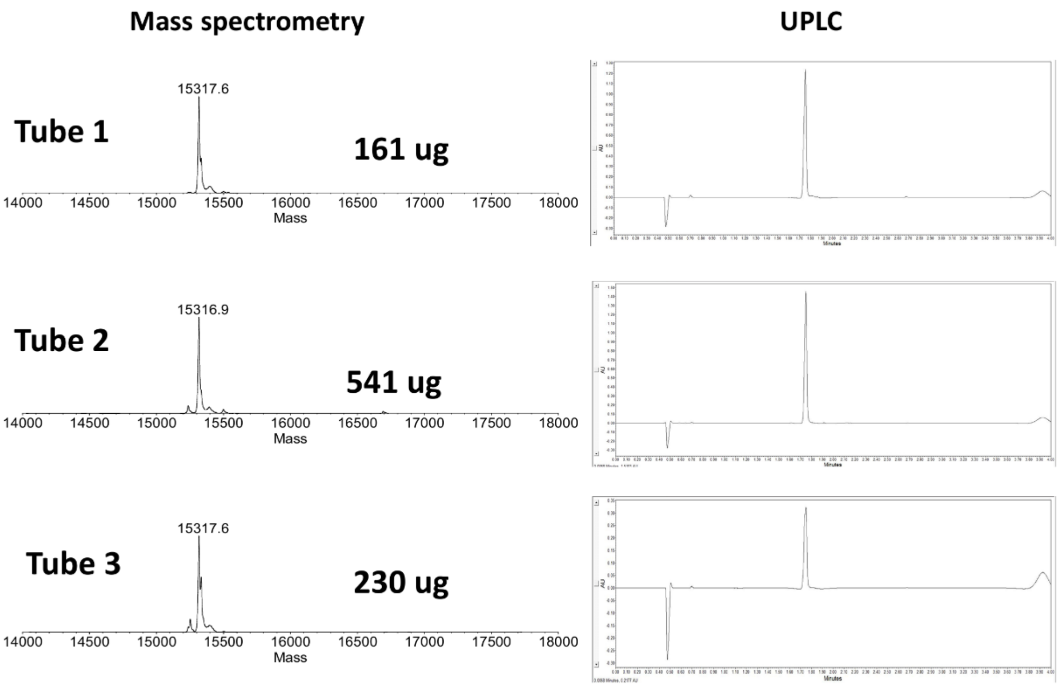


**Supplementary Fig 3 | LC-MS (ESI-LTQ) and UPLC characterization of S129-phosphorylated $^{13}C/^{15}N$ human aSyn.**
